# Functional Connectivity of the World’s Protected Areas

**DOI:** 10.1101/2021.08.16.456503

**Authors:** A. Brennan, R. Naidoo, L. Greenstreet, Z. Mehrabi, N. Ramankutty, C. Kremen

## Abstract

Rapid environmental change threatens to isolate the world’s wildlife populations and intensify biodiversity loss. Global policies have called for expanding and connecting the world’s protected areas (PAs) to curtail the crisis, yet how well PA networks currently support wildlife movement, and where connectivity conservation or restoration is most critical, have never been mapped globally. Here, we map the functional connectivity (how animals move through landscapes) of the world’s terrestrial PAs for the first time. Also, going beyond existing global connectivity indices, we quantify national PA-connectedness using an approach that meaningfully represents animal movement through anthropogenic landscapes. We find that reducing the human footprint may improve national PA-connectivity more than adding new PAs; however, both strategies are critical for improving and preserving connectivity in places where the predicted flow of animal movement is highly concentrated. We show that the majority of critical connectivity areas (CCAs) (defined as globally important areas of concentrated animal movements) remain unprotected. Of these, 72% overlap with previously-identified global conservation priority areas, while 3% of CCAs occur within moderate to heavily modified lands. Conservation and restoration of CCAs could safeguard connectivity of the world’s PAs, and dovetail with previously identified global conservation priorities.

---

Our current global system of protected areas (PAs) has been unable to slow biodiversity loss^1,2^ or provide adequate coverage for many threatened species and ecosystems^1,3–7^. PAs are constrained in size, ecological representation and governance^8^, and 90% or more exist within a matrix of human-dominated, fragmented land^9^ that is rapidly eroding^10,11^, endangering animal movement^12,13^ and survival^14^. As a result, PAs and the animal populations they contain can become isolated, interrupting the flow of vital ecological (e.g., migration, dispersal) and evolutionary (e.g., gene flow) processes that maintain populations, ecosystems and adaptive capacity^14–17^. For these reasons, the Aichi Target 11 of the Convention on Biological Diversity’s 2020 Strategic Plan for Biodiversity stipulated expanding the global PA network not only to cover 17% of terrestrial areas but also to ensure connectivity among PAs^18^, to facilitate the flow of individuals, genes and processes^19^. While these targets remained unmet by the end of 2020, post-2020 global biodiversity framework discussions continue to champion the importance of connectivity, both as a stand-alone target and integrated into other relevant targets^20,21^. To date, only a few evaluations of the connectedness of the world’s PAs exist^9,22,23^ and none explicitly map the functional connectivity of PAs. This research gap prevents the identification of globally important areas for wildlife connectivity and the spatially explicit assessment of potential threats to connectivity.

Here, we modelled the functional connectivity of terrestrial PAs (excluding Antarctica) to quantify the connectedness of each PA and map the world’s most critical areas for connectivity conservation. Our analysis focuses on the following questions relevant to global conservation practice and policy: (1) Where are the most and least functionally connected PAs in the world located? (2) How does the functional connectivity of national PA-networks compare to previous estimates of national connectivity, and what mechanisms exist for national governments to improve the functional connectivity of their PA-networks? (3) Where are the world’s most important connectivity landscapes? and (4) How do these vulnerable connectivity landscapes overlap with existing global prioritization schemes for biodiversity conservation? To answer these questions we first generated a global resistance-to-movement surface from empirical estimates of animal movement (624 individuals of 48 mammalian species) in response to the human footprint index (HFI)^12,24^, which describes the combined effects of infrastructure, land use and human access across the planet. By incorporating observed animal movements, our analysis permits a first-ever assessment of global functional connectivity, reflecting the predicted flow of mammal movement between the world’s PAs.

We applied circuit theory^25–27^ to the resistance-to-movement surface using two distinct approaches that relate the flow of animal movement across a landscape to the flow of electrical current across a circuit. First, we quantified the ‘effective resistance’^26–29^ – previously shown to predict gene flow^27,30^ – between each terrestrial PA and all other PAs on the same landmass (i.e., continent or island). Effective resistance relates the total resistance between nodes in a circuit to the degree that each node (in our case, each PA) is isolated from all others. PAs with high effective resistance are therefore further away from other PAs and/or are more difficult to travel to, based on the resistance-to-movement surface; we thus use effective resistance as an index of PA isolation (hereafter referred to as PAI). Second, we mapped the flow of electrical current across continents. For each continent we used the center grid cell of each PA as a ‘ground’ to which electrical current flowed from all other PAs^28^, proportional to PA size, with current flowing in greater density in regions of lower resistance. We summed the results from all PAs to produce a global map of relative electrical current densities that reflects the net movement probabilities of randomly moving individuals (i.e., mammal movement probability [MMP]) across all possible land-based travel routes between all terrestrial PAs. To validate our results, we compared our global map of MMP against independent GPS data from 366 individuals representing 7 species of mammals (see Methods).

## Protected Area Isolation

Does our PA isolation (PAI) index reveal new information about the connectedness of the world’s PAs? While our results agree with the expectation that the least isolated PAs occur within the world’s two most intact biomes – boreal forest and tundra^31,32^ (Figs. 1 and S1) – we nonetheless found stark contrasts between our PAI index and several global connectivity indicators previously developed for monitoring progress towards post-2020 targets^33^ (Fig. 2). Key to these contrasts is that each global indicator employs a different methodological approach to assess a different component of connectivity (Fig. S2). For example, we found that PAI, used to assess functional connectivity, varied considerably among countries that were assigned a connectivity value of zero by the ConnIntact indicator^9^, which is used to assess the contiguity of intact natural land between PAs irrespective of animal movement (i.e., structural connectivity) (Fig. 2). Indeed, PAI was only weakly correlated with all existing global connectivity indicators (Pearson’s *r* ranged from 0.36 – 0.55; Fig. 2) and it identified a different set of countries as most connected (Table S1). For example, our PAI index identified Canada, which has the second-largest area of wilderness after Russia^34^, as the 2^nd^ most connected country, while the other three indicators identified Canada as only the 15^th^, 52^nd^ or 111^th^ most connected country. Our results suggest that the PAI index we present here provides a new view of connectivity that complements how connectivity is evaluated by the existing global indicators.

**Figure 1.**
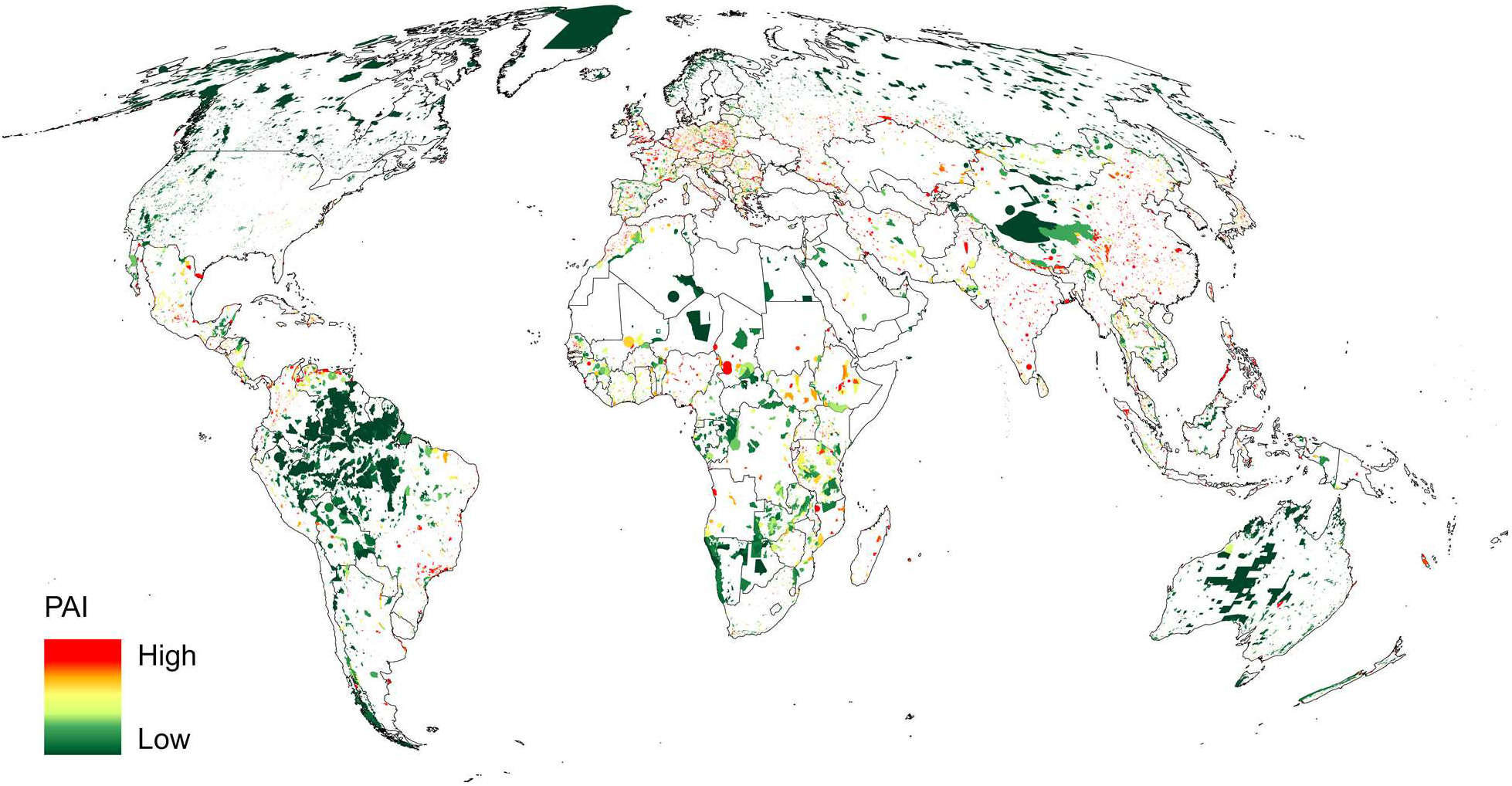
Protected area isolation (PAI). Isolation of the world’s terrestrial PAs, as measured by effective resistance to mammal movement. The most connected PAs are those categorized as wilderness areas (i.e., IUCN category Ib) and those that occur within the most intact biomes (boreal forest and tundra), while the most isolated PAs occur in mangroves (a critically endangered biome with few PAs).

**Figure 2.**
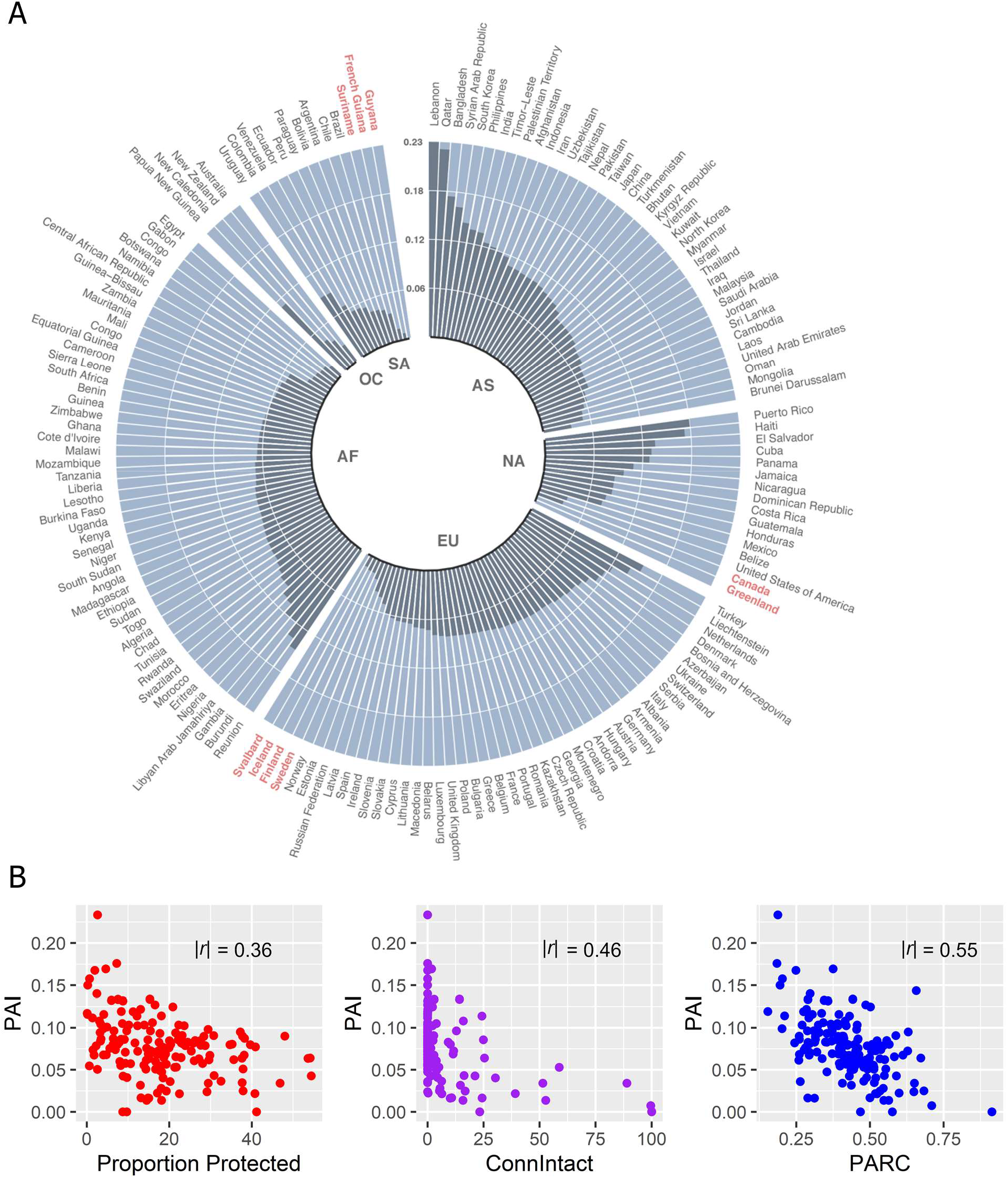
National protected area isolation (PAI). (A) PAI aggregated to the national level (dark grey bars depict PAI values); higher values reflect greater degree of isolation. Bars are organized by continent (AS = Asia; NA = North America; EU = Europe; AF = Africa; SA = South America; OC = Oceania and Australia). Red-labeled countries have the most connected national PA networks (95^th^ percentile). (B) Comparisons of our national PAI index to three existing global indicators of connectivity. ConnIntact^9^ is a recently updated version of the protected-connected index^22^; PARC is the PA connectedness index^59^. The PAI and global indicator values were weakly correlated, as evaluated using Pearson’s *r*.

What potential mechanisms exist for national governments to improve their PAI value? Because restoration^35^ and PA expansion^36^ are both recognized strategies for biodiversity conservation, we compared effects on national PAI from decreasing the human footprint (e.g., via the restoration of degraded habitats^37^) and increasing PA coverage, using a linear mixed effects model with continent as a random intercept. We found that reducing a country’s aggregate human footprint would reduce PAI more strongly than would increasing a country’s PA estate (Fig. S3). Although these strategies do not entail equal costs or effort, we found that on average a 50% reduction in the human footprint would decrease national PAI by 24% (95% CI: 17-45%), whereas a 50% increase in PA coverage would decrease national PAI by only 11% (95% CI: 9-21%). Utilizing both strategies in combination, however, has the greatest benefit, decreasing PAI by 42% (95% CI = 27-73%). These results suggest that habitat restoration, and favorable land management practices that improve the permeability of anthropogenic landscapes to animal movement^35,37,38^, can complement formal protection efforts to improve connectivity.

## Global Connectivity Conservation Priorities

Beyond the connectedness of each of the world’s PAs, can we also identify where connectivity is most vulnerable and in need of protection or restoration around the world? We focused on identifying critical concentrations of MMP that typically occur through corridors and pinch points (Fig. 3). Because these areas have few or no alternative paths for animal movement, they can have disproportionate impacts on connectivity if further restricted or destroyed^27^. We identified concentrated flows using MMP grid cells in the 90th percentile, hereafter referred to as critical connectivity areas (CCAs). We further divided CCAs into two categories based on a threshold HFI value of 4 previously used by others^9,11,31^: “modified” CCAs where HFI ≥ 4, and “intact” CCAs where HFI < 4. We found that 97% of CCAs occur in intact areas and 3% are modified (i.e., 9.7% and 0.3% of the terrestrial realm, respectively). Modified CCAs thus represent small but critical pinch points that if conserved or managed to limit further modification (e.g., through conservation easements, payments for ecosystem services, or community-based conservation^38^) could achieve gains in safeguarding connectivity through anthropogenic landscapes. The vast majority of these modified CCAs are in sub-Saharan Africa, Europe and South America, with the greatest total coverage occurring in the Bolivia, Tanzania, Democratic Republic of Congo and Poland (Fig. S4). The greatest total coverage of intact CCAs on the other hand, are in Russia, Brazil, Australia and Canada (Fig. S5).

**Figure 3.**
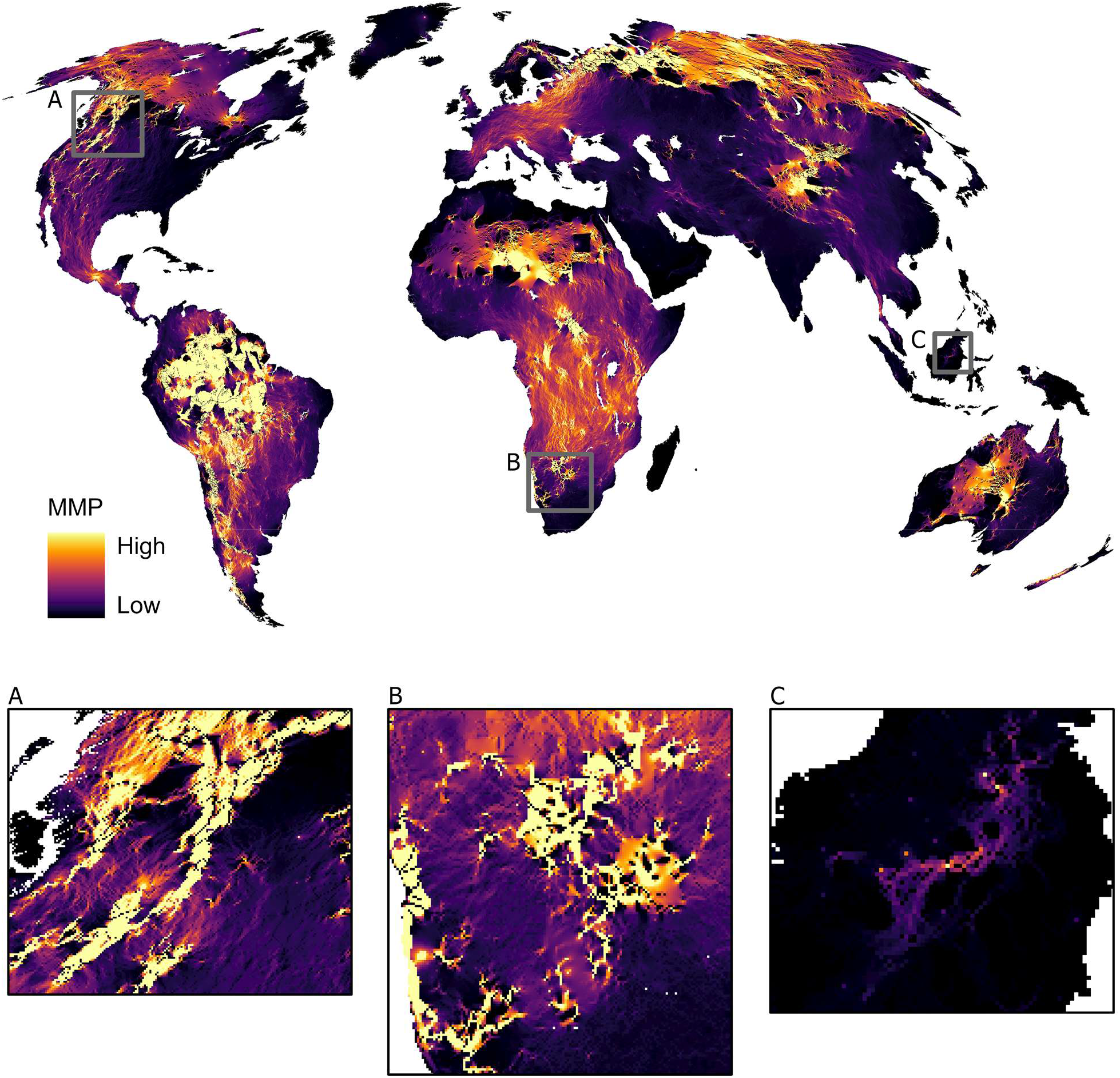
Global mammal movement probability (MMP). Predicted flow of mammal movement between all PAs based on patterns of electrical current flow. Grid cells with the highest MMP reflect areas where mammal movement is predicted to be most concentrated across the globe, flowing through corridors or pinch points (in light yellow) surrounded by less permeable anthropogenic landscape. Areas in orange and purple reflect moderate flows dispersed across many pathways. Areas of concentrated and dispersed flow are both important to connectivity, but with multiple pathways in dispersed areas, there is a lower risk of total loss of connectivity. Black regions depict areas of lower relative flow (low MMP). Boxes highlight example landscapes. Box A: corridors through mountains of western North America (e.g., Yellowstone to Yukon corridor). Box B: corridors and dispersed flow across sub-saharan Africa’s Kavango-Zambezi Transfrontier Conservation Area and coastal deserts of Namibia. Box C: flows through rainforests of Indonesia and Malaysia (e.g., Heart of Borneo conservation area).

Where are CCAs unprotected and thus vulnerable to further land-use conversion? We found that two-thirds of all CCAs are currently unprotected (Fig. 4) and that this lack of formal protection is particularly widespread for intact CCAs. We found small but important clusters of unprotected modified-CCAs in parts of Europe, sub-Saharan Africa and North and South America, where wide-ranging species, such as red deer *Cervus elaphus*, African wild dog *Lycaon pictus* and jaguar *Panthera onca*, can occur. Further, roughly 24% of all CCAs are both unprotected and occur on land suitable for future agricultural expansion^39^ (Fig. S6). The potential for rapid land conversion to agriculture highlights the need to act quickly to secure these regions for long term connectivity, either via formal protection measures or, with working lands conservation practices that focus on managing for permeable agricultural landscapes^37,38^. Such working lands, managed using strategies to enhance biodiversity conservation (e.g., silvo-pastoral, agroforestry and other agroecological management practices) and benefits to humans (e.g., pollination services and pest control), could also represent an important other effective area-based conservation measure (OECM) and as such could provide important contributions to global conservation policy targets^18,21^.

**Figure 4.**
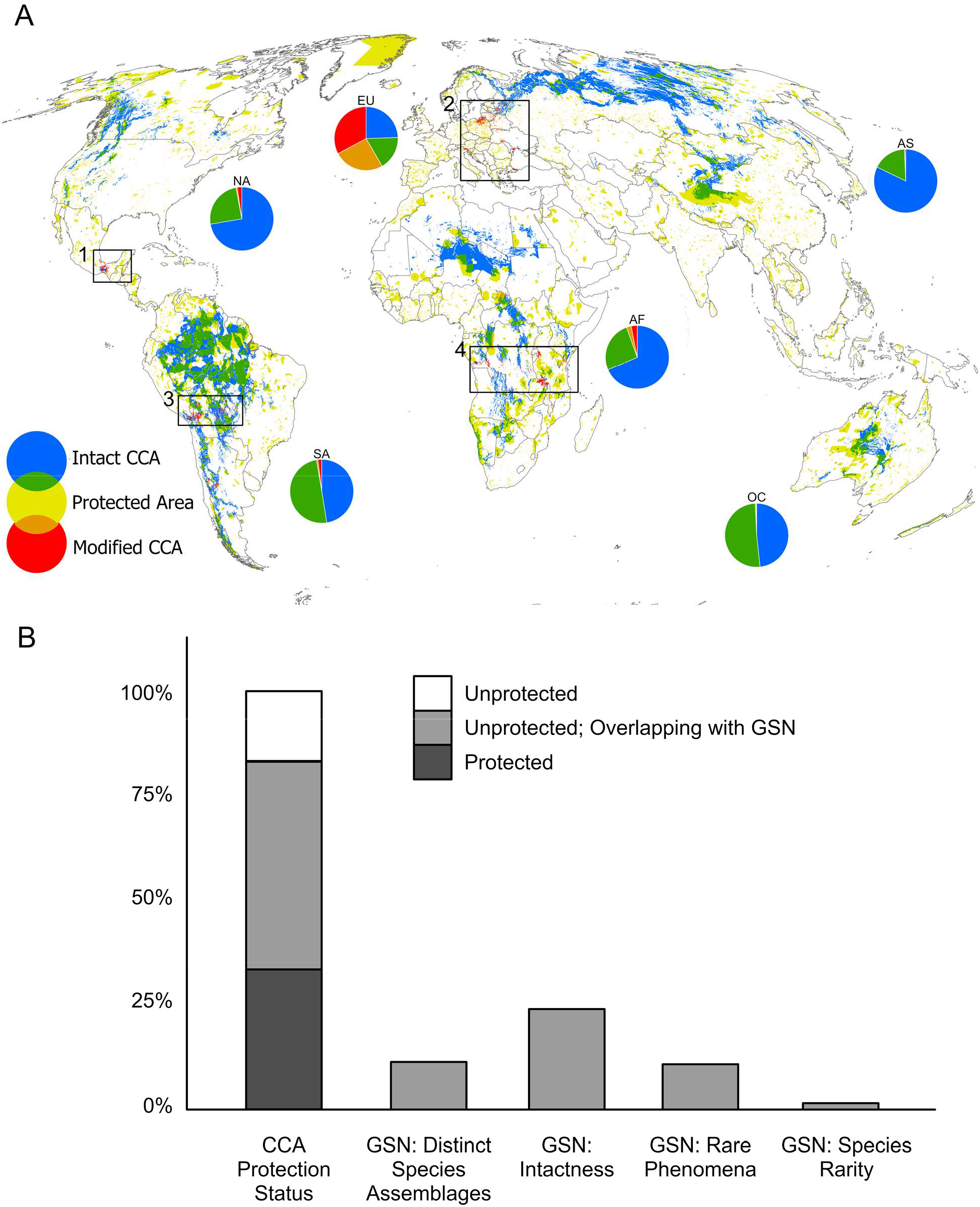
Mapping critical connectivity areas globally (CCAs). (A) Intact CCAs (blue) and modified CCAs (red) were classified based on mammal movement probability (MMP) and human footprint index values (HFI< 4, HFI ≥ 4, respectively). CCAs in green and orange are covered by protected areas. Black boxes highlight four areas with clusters of modified CCAs, occurring in: (1) Mexico, (2) Poland, Slovenia, Italy, Belarus and Ukraine, (3) Bolivia and Brazil, and (4) Democratic Republic of Congo, Tanzania and Zambia. Pie charts indicate the proportion of each CCA type and protection status in each continent (AS = Asia; NA = North America; EU = Europe [excluding Russia]; AF = Africa; SA = South America; OC = Oceania and Australia) (B) Proportion of CCAs that overlap with expanded conservation areas of the Global Safety Net (GSN). Stacked bars (far left) depict the proportion of protected and unprotected CCAs. Individual bars (right) represent the proportion of CCAs that overlap with each of the priority conservation areas selected for specific purposes under the GSN.

Which of the world’s continents, biomes and ecoregions contain the greatest proportion of CCAs? We found that the tropical moist forests of South America, the tropical grasslands and deserts of Africa, and the boreal forests of Asia contain > 80% more CCAs than other biomes (Fig. 4). Globally, these four biomes contain 41% of the world’s ecoregions and a majority of the world’s species, 50% of which are contained within tropical moist forests^40^. Of the world’s 846 ecoregions, the 50 that were recently identified as having the greatest potential to protect biodiversity^36^ also contribute disproportionately to connectivity; unprotected portions of these priority ecoregions contain on average two times greater MMP than all other unprotected ecoregions (Fig. S7) and contain > 42% of the unprotected CCAs. Thus, conserving movement corridors and restoring degraded habitat around pinch points in the unprotected portions of these priority ecoregions is likely to benefit biodiversity conservation *and* safeguard the connectivity of currently established PAs, a key goal in the post-2020 global biodiversity framework^21^.

## Connectivity-Biodiversity Conservation Synergies

Where can we further maximize conservation effectiveness by identifying landscapes that simultaneously enhance connectivity and biodiversity? We examined the proportion of CCAs that overlap with the Global Safety Net (GSN) – a proposed global conservation scheme that identifies new priority areas for expanded protection that would boost the world’s terrestrial PA coverage to ∼ 50%^36^. We found that roughly 72% of unprotected CCAs, and a majority of CCAs suitable for future agricultural expansion, overlap with the GSN priority areas for expanded conservation (Figs. 4 and S7). Thus, the GSN would protect key biodiversity elements as well as more than 81% of the world’s most important areas for connectivity. Of the remaining CCAs not covered by the GSN, many are vulnerable to potential disturbance, degradation and/or land-conversion (9% are modified and roughly 30% have the potential for agricultural development; Fig. S7).

We also examined CCA overlap with other global conservation prioritization schemes that were not explicitly incorporated into the GSN, including Frontier Forests^41^, Global 200 Ecoregions^42^ and High-Biodiversity Wilderness Areas^43^. We found that > 60% of the CCAs overlap with currently-unprotected portions of these global conservation schemes, and that the Global 200 Ecoregions covered the highest percentage of CCAs (∼30%; Fig. S8). This overlap between CCAs and global conservation priorities highlights potential synergies that conservation efforts in these areas could deliver via maintaining globally-significant areas for connectivity, while also preserving rare species and assemblages, biologically outstanding ecoregions and intact habitats critical for biodiversity and ecosystem function.

## Conclusions

Our study identifies key opportunities for conserving globally important areas for connectivity and for monitoring national PA connectivity using circuit theory methods adapted to the global scale. Our results show that the functional connectivity approach substantially changes the view of national PA connectivity compared to the existing global-policy indicators that are more geared towards assessing other aspects of connectivity. A key difference is that our approach specifically incorporates observed animal responses to attributes of natural and human-dominated landscapes, which is lacking in the other global indicators. Our global application of circuit theory also makes it possible for the first time to assess the permeability of both intact and human-modified landscapes to mammal movement across the terrestrial realm. With this approach, we identified small areas of modified but still permeable lands, representing only 0.3% of the terrestrial world, where timely conservation is of utmost importance to prevent further impediments to the flow of mammals between PAs.

Our study specifically evaluates the functional connectivity of protected areas at the global scale, and therefore we do not explicitly consider the connectedness of other potential sources and destinations of movement (e.g., OECMs or other natural areas), nor do we portray connectivity for more localized movements (e.g., inter-patch movement). In addition, our connectivity model is informed only by mammal movements, and therefore may not capture connectivity for other taxa. Indeed, our results illustrate that differences in national PA connectivity can occur even among mammalian species; we found that PAs appear more isolated from the perspective of larger mammals (see Methods and Fig. S9). While this suggests that larger mammals are more sensitive to higher HFI levels, we found that patterns of flow – that is, where the flow of mammal movement is dispersed versus constricted – remain consistent across mammalian body sizes. Finally, although our approach is based on mammalian movement, our results illustrate how CCAs spatially overlap with global conservation priorities that aim to protect a variety of taxonomic groups.

We found that the vast majority of the world’s CCAs currently under threat (i.e., unprotected or located on modified land), or likely to be threatened in the future (i.e., located on land suitable for future food production), would be protected by expanded conservation under the proposed Global Safety Net (GSN)^36^. If protected, these areas of GSN-CCA overlap could result in important conservation synergies that slow or prevent biodiversity loss. Since many of these areas of overlap are also predicted to be suitable for future agricultural expansion, where formal protections could be contested over livelihoods or food supply needs, alternative strategies for conserving or restoring connectivity may be needed. Our results suggest that managing the matrix for more permeability to animal movement (e.g., habitat restoration) can be a better single strategy for improving connectivity than adding protected areas alone. However, formal protection and habitat restoration together generated the largest improvements to connectivity, supporting the need for holistic approaches that combine working lands conservation^37,38^ with protected areas. Such combined strategies can also provide significant benefits to humans^37,38,44^, thus advancing the post-2020 global biodiversity framework vision of “living in harmony with nature”^21^.

## Methods

We modelled connectivity using Circuitscape, an open-source program implemented in Julia^45^ and widely used in research and conservation planning^25,27,46–51^ that relies on principles from circuit theory to relate the flow of animal movement across a heterogeneous landscape to the flow of electrical current across a circuit^27^. This approach treats a gridded resistance-to-movement surface as a conductive layer where neighboring grid cells are connected by resistors, whose level of resistance represents the friction of the landscape to animal movement (i.e., low-resistance grid cells are most likely to be traversed). Electrical current, injected at a source node representing a source of animal movement, flows across all neighboring landscape grid cells (resistors) in the circuit to a ground node representing the destination of animal movement. Connectivity is then evaluated in two distinct ways: (1) from the effective resistance of the circuit, measuring the degree to which the source node is isolated from the ground node and (2) from the pattern of electrical current flow (i.e., animal movement probabilities) across the resistance-to-movement surface^27^, as outlined below and illustrated as a workflow in Fig. S10.

Two inputs are required for this analysis: the resistance-to-movement surface describing the friction of the landscape to animal movement and the layer specifying sources and destinations of animal movement. To model functional connectivity (Fig. S3), we generated a resistance-to-movement surface using the data and linear mixed effects model of 0.95 quantile displacement distances from 10-day intervals presented in Tucker et al. (2018)^12^ for 624 GPS-collared individuals from 48 species of mammals. This movement model included as predictor variables the human footprint index (HFI)^24^, normalized difference vegetation index (NDVI), body weight and dietary guild, and species as a random effect. We focused on the long-distance, long-time scale movements as those more likely to represent the scale of movement occurring between protected areas, thus representing movements promoting gene flow, rescue effects or climate adaptation.

Using the model above, we predicted mammal movement for all values of HFI across Earth’s terrestrial surface after resampling the HFI layer from 1 km^2^ to ∼70 km^2^ (in Mollweide equal area projection) to reduce computation time in our Circuitscape analysis from months to days (using a Google virtual machine with 96 CPUs and 360 GB memory). We rescaled these predicted movements to between 0 and 1 to represent the landscape’s permeability to mammal movement across the globe. We then used an inverse linear function (e.g., 1-permeability) to convert the permeability surface to a resistance-to-movement surface^52–54^. Following other studies, we further modified resistance-to-movement, M, by percent slope to reflect the energetic costs of moving in steep terrain^28,50,55^ using the equation 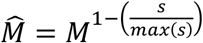, where s = terrain slope.

We used all terrestrial PAs from the May 2020 World Database on Protected Areas (WDPA)^56^ as the sources and destinations of mammal movement (i.e., source and ground of electrical current). We combined this WDPA layer with PAs in China from the April 2018 WDPA^56^, because China removed all its national PAs from the WDPA after this date. As recommended in the WDPA best practices guide, we removed any PA that did not report its area, or with a ‘Proposed’ status or ‘UNESCO-MAB Biosphere Reserve’ designation (https://www.protectedplanet.net/en/resources/calculating-protected-area-coverage). We used PA centroids as the source and ground of electrical current, rather than entire PA polygons, because this allowed for estimates of electrical current flow within PAs, rather than assuming flow is equally good within PA boundaries^28^. We rasterized PA-centroids to match the resolution and extent of the resistance-to-movement surface. We excluded PA-centroid grid cells for which the sum total of PA area was less than half the grid cell resolution, 35 km^2^, to ensure that electrical source grid cells are primarily represented by a probable source or destination of mammal movement. By excluding these small PAs (similar to^28,57^), we also avoided overestimating connectivity from using source and destination nodes much smaller than the resolution of our analysis and the home ranges of many of the mammals that we examined. Excluded PAs represented 90% of the total number of individual PAs but, with a mean area of 2.4 km^2^, they comprised only ∼2% of the total area of the global PA estate (Fig S11 and Table S2). Included PAs were on average 420 times larger than excluded PAs. Note also that excluded PA-centroids were removed only as sources and destinations of mammal movement in the model; importantly their benefits to connectivity were fully incorporated into the resistance-to-movement surface as areas of lower resistance (i.e., lower HFI). We modelled connectivity by continent, excluding Antarctica, islands < 2,500 km^2^ and islands with < 2 PAs. We also combined Europe and Asia to avoid boundary effects where the two continents meet.

### Protected Area Isolation

To obtain a value of effective resistance for all terrestrial PAs (steps 4-5 in Fig. S10), we applied Circuitscape’s one-to-all mode^58^ to the input layers. This mode connects a focal PA-centroid to an electrical current source and connects all remaining PA-centroids to ground, thereby generating one circuit where electrical current flows from the focal PA source to all other PAs within the continent. When the circuit is connected, effective resistance, R_eff_, is calculated as 1/R_eff_ = 1/R_1_ + 1/R_2_ +1/R_3_ … 1/R_n_, where each R_i_ represents a pathway (i.e., summed grid cell ‘resistors’ connected in series) between PAs. Following this central equation of circuit theory, effective resistance decreases, and thus connectivity increases, as more connections (i.e., redundant pathways, R_i_) are added to the circuit^27^. For this process we maintain the default current injection of 1 Amp, because each grid cell ‘resistor’ in the landscape has an assigned resistance value, effective resistance does not vary with the amount of current injected. We repeated this procedure for each PA and mapped the resulting PA isolation (PAI) values for all qualified terrestrial PAs (i.e., with summed area within a grid cell of > 35 km^2^) and evaluated PAI within the world’s biomes^42^ and continents.

We calculated the median PAI for each country’s PA network. Because measures of PAI were generated from connecting PAs across entire continents, national PAI may be affected by connections with neighboring cross-border PAs. To ensure that national PAI was not unduly sensitive to the location and extent of PAs outside the country’s borders, we repeated the steps to model PAI at the country level for 25 randomly selected countries. To do so, we clipped the resistance-to-movement surface and PA layer to each of the 25 countries, re-applied circuit theory using the one-to-all approach and re-computed median PAI by country. National PAI from the continent and country analyses were highly correlated (Pearson’s *r* = 0.92; Fig. S12), suggesting that metrics from our global analysis are suitable proxies for country-level PAI.

We evaluated how hypothetical changes to mean-national HFI and proportion of PA would affect national PAI. We used a linear mixed effects model with mean HFI and % PA area as predictor variables scaled by subtracting the mean and dividing by the standard deviation to evaluate the relative effects of each, with continent as a random intercept. We then used the estimated coefficients to simulate the effects of changing the proportion of protected area (including PAs of all sizes) and mean HFI on national PAI.

We compared our national PAI index, which explicitly accounts for mammal movement, to three proposed indicators of progress towards post-2020 global connectivity targets that lack measurements of animal movement (Fig. S3): (1) the proportion of country protected, (2) ConnIntact^9^ – an updated version of Protconn^22^, the currently proposed global indicator – and (3) the PA connectedness index (PARC)^59^. ConnIntact is a network-based structural connectivity approach that accounts for the number of PAs, total area of PAs, total country area and the contiguity of intact land between PAs. Specifically, this approach assumes PAs are perfectly connected whenever there is a contiguous pathway of 1 km^2^ grid cells of intact natural land directly connecting PA edges or indirectly connecting PAs through connecting intermediate PAs as stepping stones. PAs are assumed to be completely disconnected when there is no contiguous pathway of intact land. The PARC index accounts for connectivity between ∼1 km^2^ grid PA grid cells and between these PA grid cells and ∼1 km^2^ grid cells containing primary vegetation using a cost-distance approach where the cost of moving between grid cells is assumed to be negatively related to the proportion of primary vegetation (i.e., the more primary vegetation in a grid cell, the lower the cost). We used Pearson’s *r* to evaluate the correlations between national PAI and the three global indicators of connectivity.

### Mapping Global Connectivity

To map patterns of mammal movement probability (MMP) using electrical current density (steps 6-10 in Fig. S10) we applied Circuitscape’s all-to-one mode^58^ to the input layers. Similar to the pairwise mode^58^, the all-to-one mode produces patterns of electrical current flow between each source-ground pair but with greater computational efficiency. It does this by generating many circuits simultaneously; in our application, one PA-centroid is designated as ground while all remaining PA-centroids are current sources. At each source, we injected an electrical current amount proportional to PA area (e.g., we injected 100 Amps into any PA centroid that represented a 100-km^2^ PA), to account for the effect of PA size on the number of animals potentially moving from each PA (i.e., assuming that all else equal, large PAs have a greater abundance of wildlife and thus more dispersing animals than small PAs^60^). To account for the size of the destination PA (i.e., all else equal, large PAs will receive more immigrating animals than small PAs^60^), we multiplied the output electrical current density map – depicting electrical current flow from all source PAs to one ground PA – by the area of the corresponding ground PA. We repeated this process for each PA-centroid connected to ground and summed all maps to obtain a map of cumulative electrical current density (i.e., MMP) per continent.

We found that electrical current densities increased with land mass area. To correct for this effect and ‘normalize’ electrical current densities into a common scale – using current density/km^2^ – we divided electrical current densities by the area of their corresponding land mass. Finally, we merged all electrical current density/km^2^ maps together to generate a comparable global map.

### Validation and Sensitivity Analyses

We validated our global map of electrical current density (MMP) using two different approaches. First, we used independent GPS movement data (> 1.5 million locations) from 366 individuals and 7 mammal species, combined with GPS movement data (> 220,000 locations) from another 88 individuals and 4 species included in the Tucker et al. (2018)^12^ analysis, to evaluate whether connectivity values were higher where animals were observed moving (see Table S3 for details and list of data providers; see Fig. S13 for dataset locations). We obtained known range data for each species (https://www.iucnredlist.org/). Then for each species-dataset we (1) generated a 95% utilization distribution (UD) (i.e., 95% kernel home range) from the GPS data using the adehabitatHR package^61^ in the R statistical computing environment^62^ version 3.6.3, (2) randomly selected 1000 points within that UD to represent locations where those individuals were likely to occur and (3) extracted electrical current density values for those points. To compare these values to electrical current density values where the collared individuals could have been but were not observed, we (4) calculated max displacement distance using the adehabitatHR package^61^ in R and buffered the UD by this distance, (5) clipped the species range data to that buffer, (6) randomly selected points within that clipped range outside of the UD and (7) extracted electrical current density values from those random points.

We used a Bayesian hierarchical approach to evaluate the differences in electrical current density extracted from points inside and outside the UDs. We used the package brms^63^ in R to construct the model, with log electrical current density as the response variable, inside or outside UD as a categorical predictor variable, and a random intercept and slope by species. We used uninformative prior distributions, ran 4 iterations of the model with 400 chains per iteration, and confirmed that all parameters converged with R-hat values < 1.1. Using the predicted posterior distributions, we calculated the percentage that electrical current densities were higher inside the UDs and provide strong evidence for species moving within areas of higher predicted movement probability (Fig. S14).

Second, using the same data, we evaluated whether animals were selecting for critical connectivity areas (CCAs) (i.e., grid cells having electrical current density values in the 90th percentile). Using a raster of CCAs and the GPS datasets described above, we extracted the number of unique CCA grid cells that were used by each species. We calculated the proportion of used grid cells that were CCAs. We also calculated the proportion of unused grid cells that were CCAs, within the same buffered areas described previously. From this, we calculated the percent that CCAs were used (i.e., inside UDs) relative to what was available and found strong evidence that 9 of the 11 species preferred to move within CCAs (Table S4).

We also evaluated the sensitivity of effective resistance (PAI) and electrical current density (MMP) to species body size^12,64^ and continent-specific species attributes. We first created three different resistance-to-movement surfaces using a similar linear mixed effects model that was used in Tucker et al. (2018)^12^ and our previous analysis, but instead fitted to: (1) 10-day displacement distances recorded by species whose mean body mass was greater than the mean body mass, ∼ 53 kg, of the 48 species considered in our full model, (2) 10-day displacement distances recorded by species whose mean body mass was less than the mean body mass of the 48 species, and (3) 10-day displacement distances recorded by African species only. We repeated the steps in the workflow described for our main analysis (Fig. S10) and compared effective resistances and electrical current densities between our full model and the sensitivity test models (Fig. S9). Effective resistances increased with body size and electrical current densities were highly correlated across models (R^2^ > 0.9), suggesting that while larger mammals are more sensitive to high HFI, the patterns of flow remain consistent across mammalian body sizes.

### Areas of Overlap for Priority Connectivity and Biodiversity Sites

We summarized the total area of both modified and intact categories of critical connectivity areas (CCAs) by country and identified countries with the largest area (95th percentile) of CCAs. To assess the protection status of CCAs, we quantified CCA overlap with PAs using the May 2020 World Database on Protected Areas (WDPA) and the April 2018 WDPA for PAs in China^56^. We used both WDPA polygons and WDPA points files to generate our PA layer. We converted WDPA points to polygons using circular buffers with areas equal to their corresponding reported PA area (https://www.protectedplanet.net/en/resources/calculating-protected-area-coverage), and then we spatially intersected these buffers with their country of origin to exclude parts of the buffer that occurred in other countries. We compared MMP in protected and unprotected portions of the 50 ecoregions recently identified as contributing most to enhancing biodiversity protection^36^ and in protected and unprotected portions of all other ecoregions (https://ecoregions2017.appspot.com/). We also summarized the total area of CCAs within continents, ecoregions^40^ and biomes^42^.

We quantified the amount of unprotected CCAs that overlapped with lands where future agricultural expansion is predicted using a global land-systems (GLS) map describing crop suitability as determined by agro-climatic and agro-edaphic conditions^39^. The crop suitable areas in the GLS map included those that were categorized as very high to marginally suitable for high intensity cropland along with low intensity cropland, and thus may overpredict cropland expansion in some cases^39^. However, suitable areas for tree plantation and livestock expansion were not included in the GLS map, potentially resulting in an underprediction of CCA overlap with future agricultural areas.

We examined the proportion of unprotected CCAs that overlapped with four areas prioritized for expanded conservation under the Global Safety Net (GSN)^36^, the most recent global prioritization scheme for biodiversity conservation. These areas are prioritized in the GSN because of their disproportionate contribution to (1) important rare and threatened species, (2) distinct species assemblages, (3) intact assemblages of large mammals and (4) wilderness or other macrorefugia for wildlife. Conservation of these additional four priority areas, plus climate stabilization sites, would extend coverage of the PA estate to ∼50% of the terrestrial realm.

We also quantified the proportion of unprotected CCAs that overlapped with three other global conservation prioritization schemes that target other critical ecological conditions or regions important to biodiversity (which may be partially overlapping with the GSN), including Frontier Forests^41^, Global 200 Ecoregions^65^ and High-Biodiversity Wilderness Areas^43^. Frontier forests represent large contiguous wilderness areas that specifically encompass forests and conditions important to forest integrity. The Global 200 Ecoregions is based on representation and biologically outstanding character, and is thus meant to encompass the most biologically outstanding ecoregions in every biome-realm combination. High Biodiversity Wilderness Areas are the complement of Biodiversity Hotspots, thus encompassing non-threatened areas of high biodiversity.

## Acknowledgements

We are grateful to Christina Kennedy for her thoughtful comments on the manuscript.

