## Supplemental Materials for "Functional Connectivity of the World’s Protected Areas"

**This file includes:**

Figs. S1 to S14
Tables S1 to S4
References

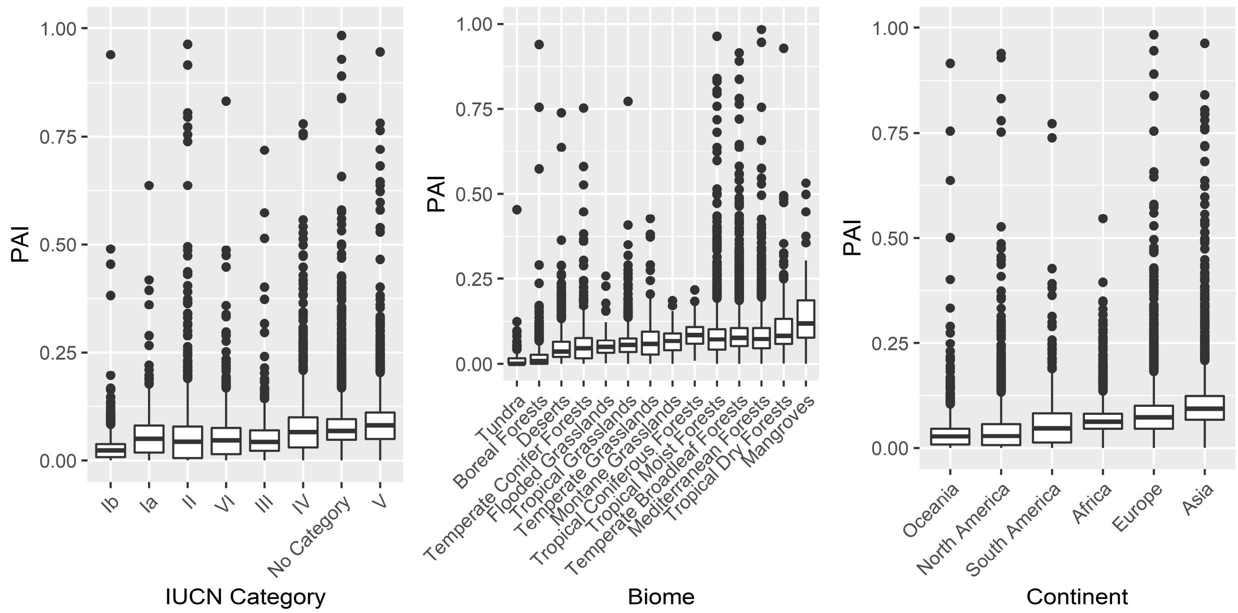


Fig. S1. Summary of protected area isolation (PAI) values. Boxplots of global PAI by PA-IUCN category (left), biome (middle) and continent (right). The least isolated PAs occur among PAs categorized as wilderness area (IUCN category Ib), within the boreal forest and tundra biomes, and across the continents of Oceania and North America. The most isolated PAs occur among PAs categorized as protected landscapes (IUCN category V), within the mangrove biome, which is highly endangered with no remaining globally significant wilderness and the fewest total number of PAs, and across the Asian continent.

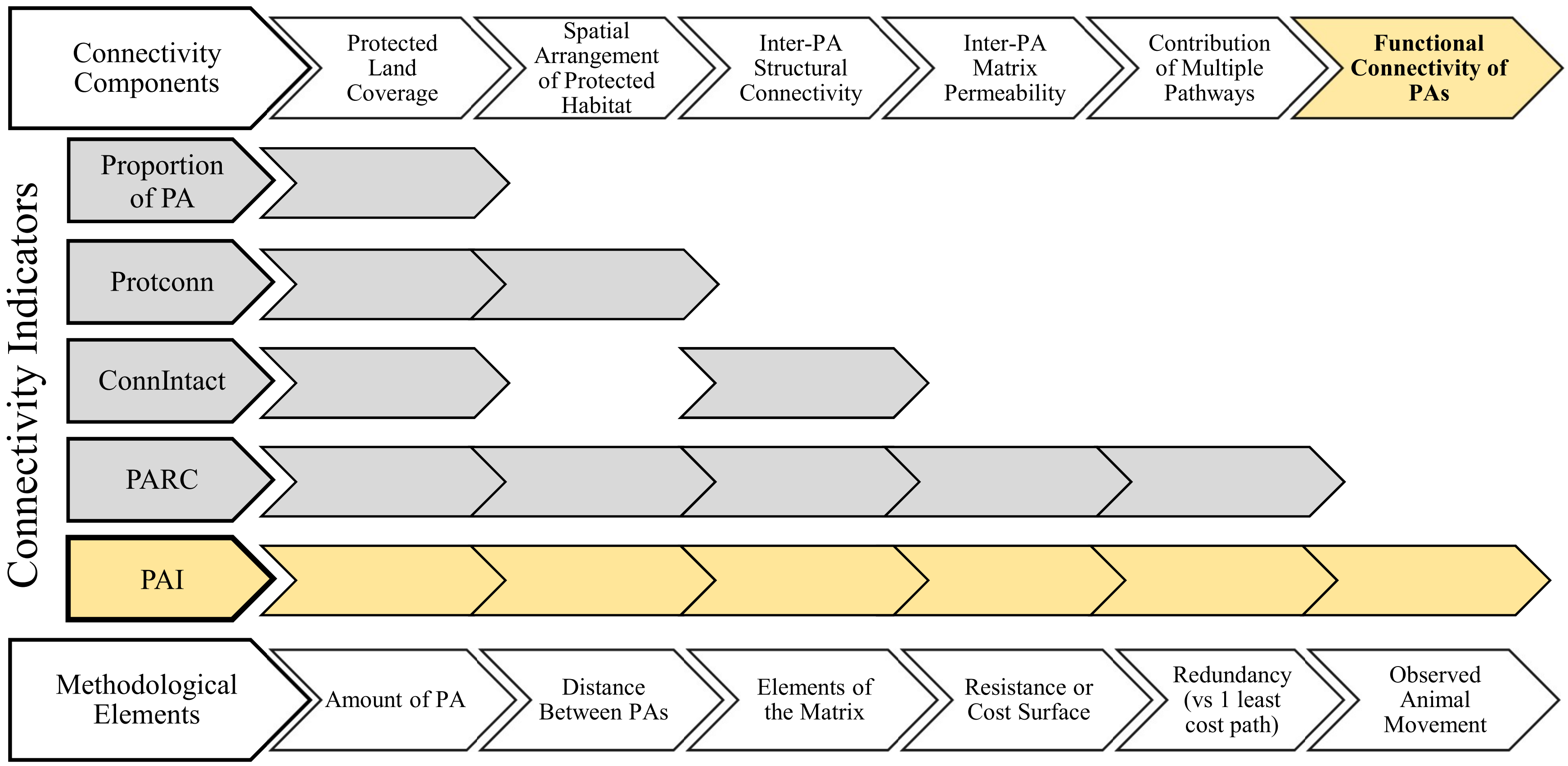


Fig. S2. The spectrum of connectivity components evaluated by existing global indicators of connectivity. Global indicators include Protconn[1] and its update ConnIntact[2], PARC[3] and our protected area isolation (PAI) index in yellow. Colored chevrons indicate the aspect of connectivity and model elements that each indicator comprises. The spectrum moves from the spatial coverage of PA to the structural connectedness of the landscape – referring strictly to the spatial arrangement of PAs and landscape elements, irrespective of how animals move – to the functional connectedness of the landscape for animal movement. Including measurements of animal movement (or gene flow) is the key attribute to evaluating functional connectivity.

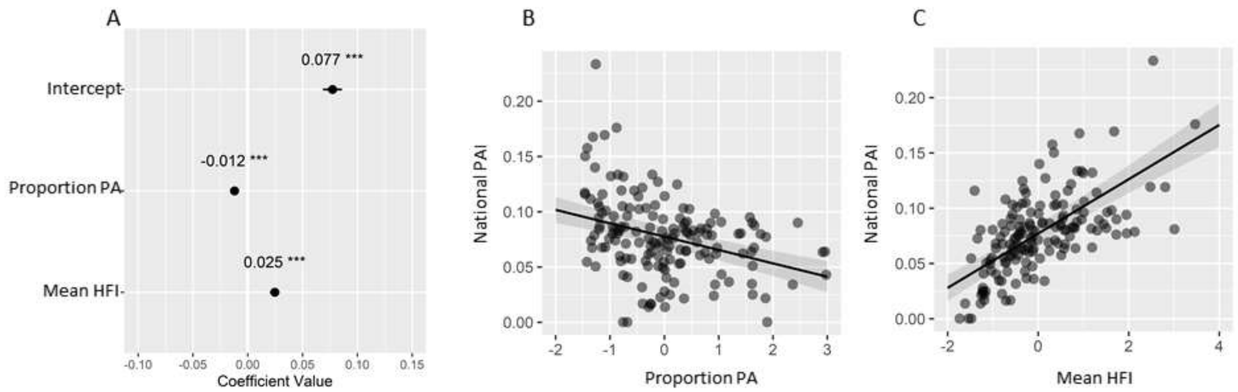


Fig. S3. Predictors of national protected area isolation (PAI). (A) Coefficient and 95% CIs estimated from a linear mixed effects model of national PAI (n = 160) with scaled predictor variables: proportion of PA (Proportion PA) and mean Human Footprint Index (HFI). CIs that are not visible fall within the dot indicating the mean value. (B) Estimated relationship between national PAI and Proportion PA. (C) Estimated relationship between national PAI and mean HFI.

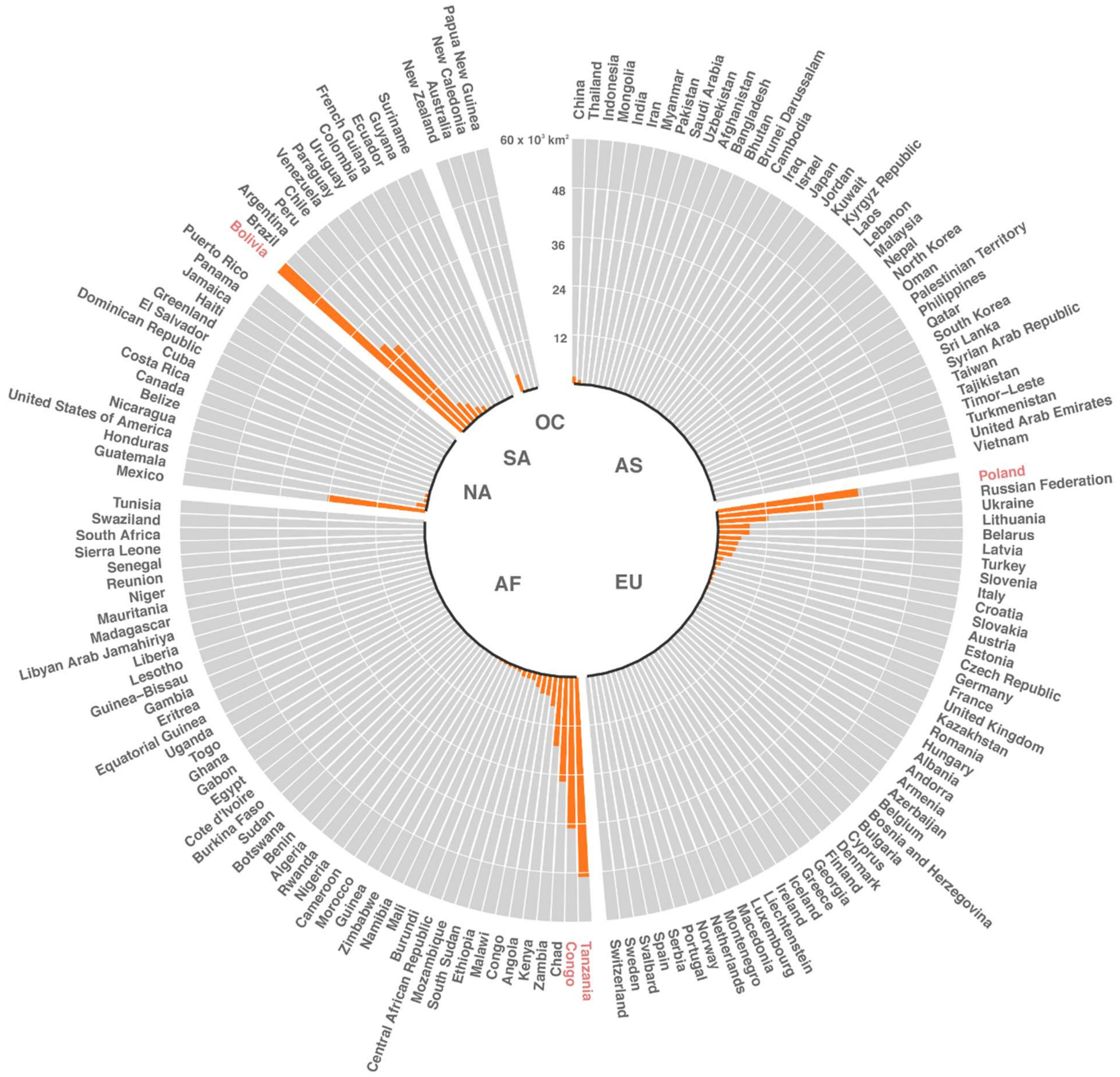


Fig. S4. National coverage (km$^2$) of modified critical connectivity areas (CCAs). Areal coverage of CCAs is organized from highest to lowest within each continent (AS = Asia; NA = North America; EU = Europe; AF = Africa; SA = South America; OC = Oceania and Australia). Red labeled countries represent countries with the most area covered by modified CCAs (upper 5%).

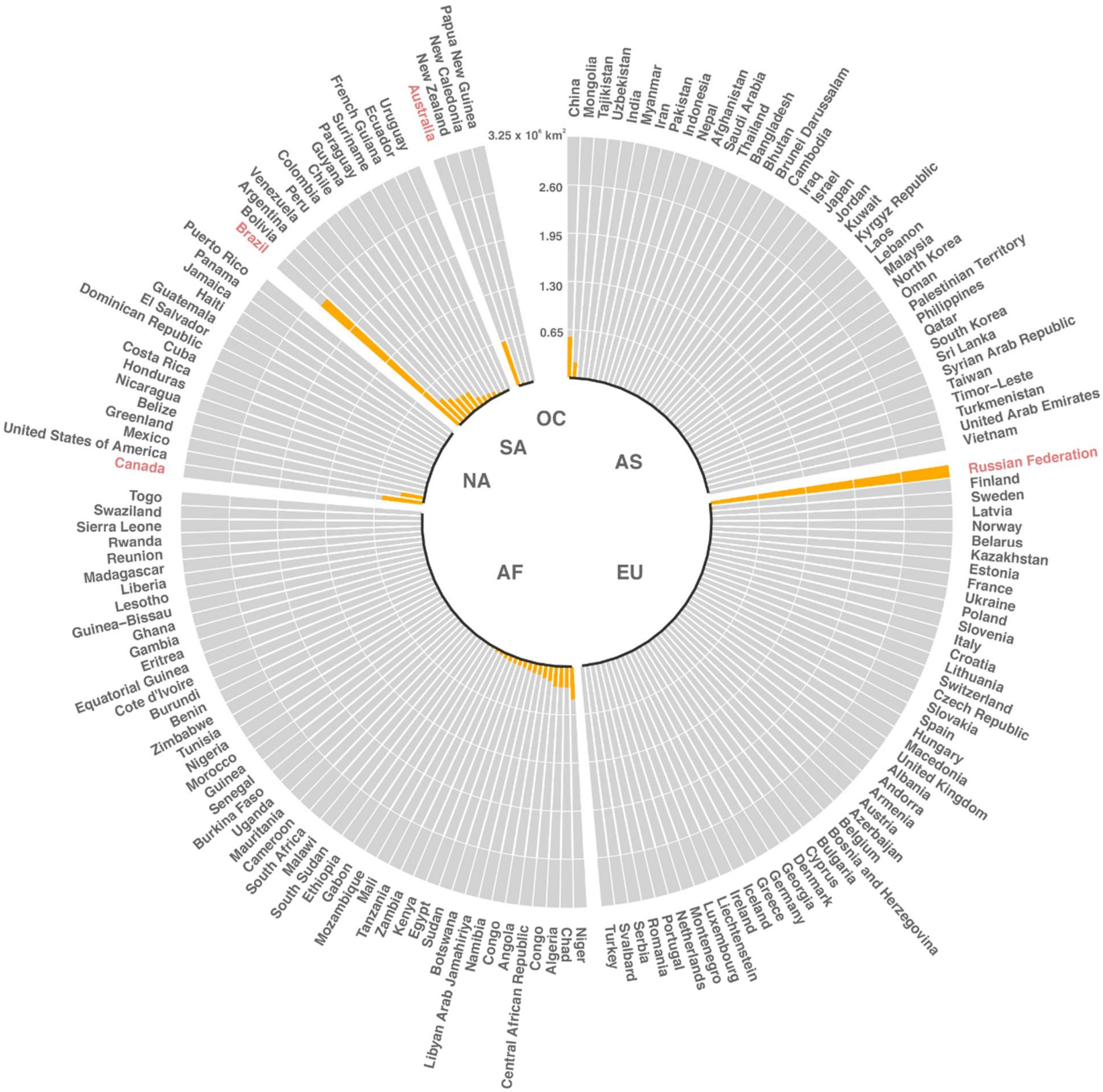


Fig. S5. National coverage (km$^2$) of intact critical connectivity areas (CCAs). Areal coverage of CCAs is organized from highest to lowest within each continent (AS = Asia; NA = North America; EU = Europe; AF = Africa; SA = South America; OC = Oceania and Australia). Red labeled countries represent countries with the most area covered by intact CCAs (upper 5%).

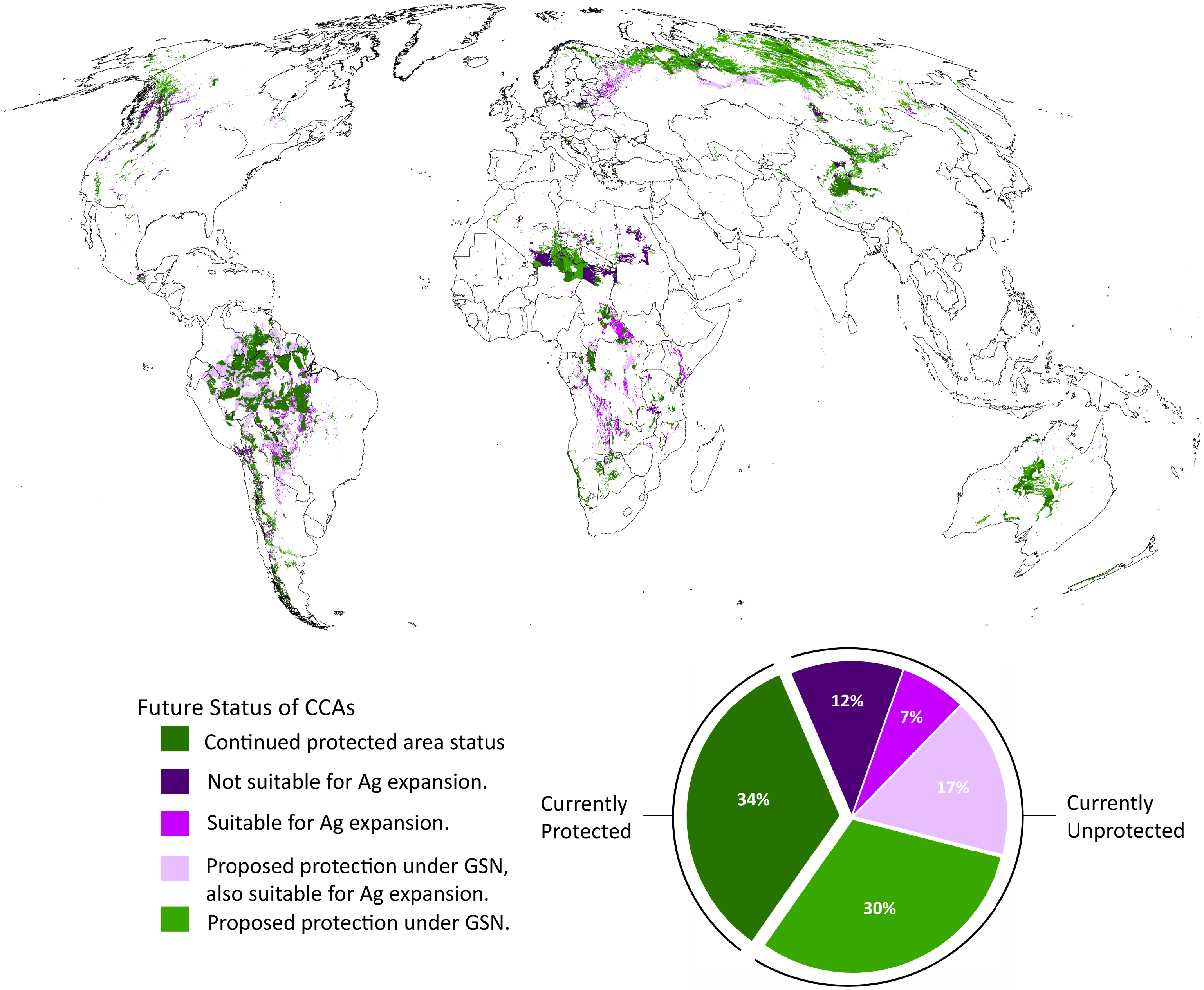


Fig. S6. Future protection and threat status of CCAs. Potential future protection occurs where currently unprotected CCAs overlap with areas prioritized for expanded conservation under the Global Safety Net (GSN). Future threats were examined where CCAs do not overlap with priority areas of the GSN (i.e., remain unprotected) and where they overlap with areas predicted to be suitable for future agricultural (Ag) expansion.

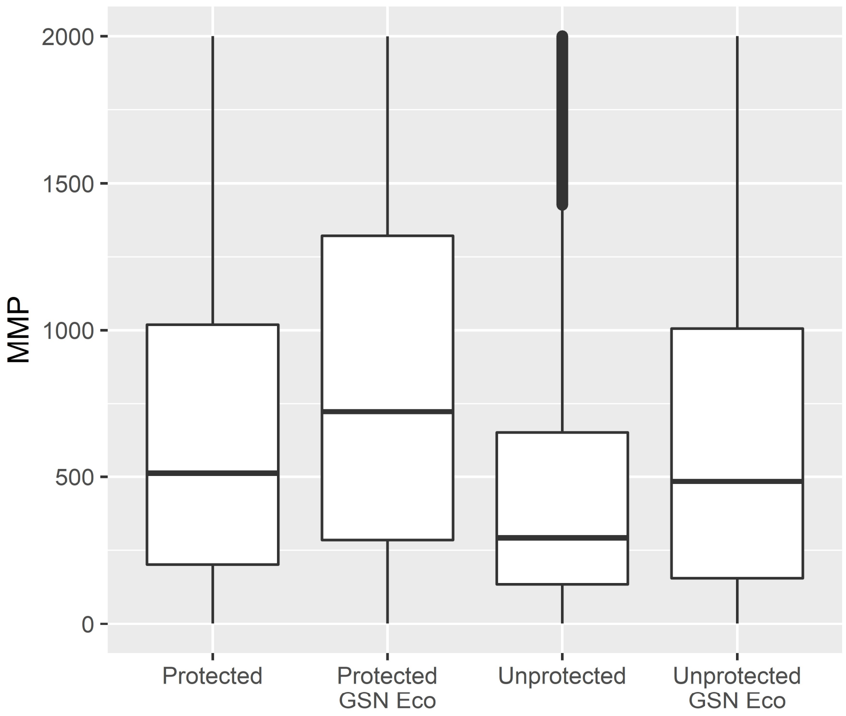


Fig. S7. Connectivity of the protected and unprotected portions of ecoregions. Boxplots of predicted mammal movement probability (MMP) within the 50 priority ecoregions identified by the Global Safety Net (GSN Eco) and within all other ecoregions, separated by protection status (Protected and Unprotected).

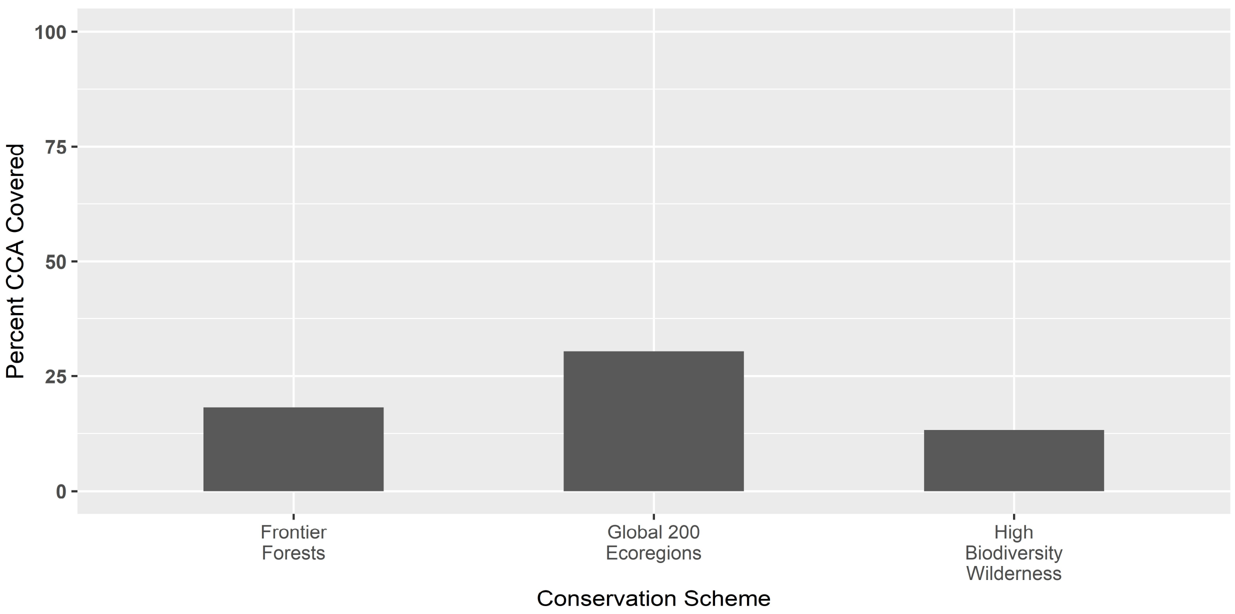


Fig. S8. Critical connectivity area (CCA) overlap with other global conservation schemes.

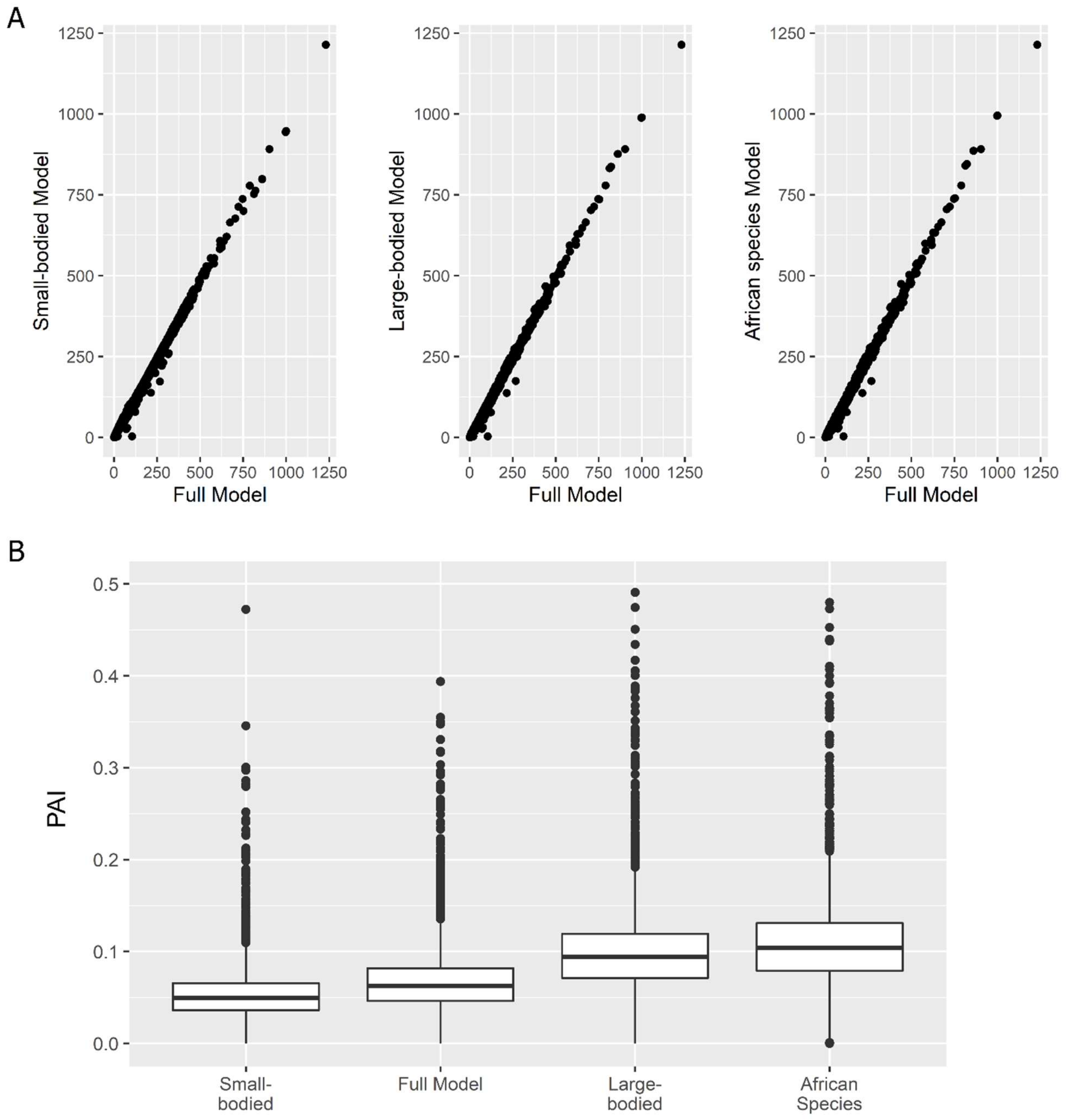


Fig. S9. Sensitivity analysis. Comparison of (A) predicted mammal movement probability (MMP) and (B) protected area isolation (PAI) between the full model in our main analysis and three other connectivity models generated using resistance-to-movement surfaces parameterized for small-bodied species in the Tucker et al. 2018[4] dataset (mean body mass = ~13 kg), large-bodied species in the dataset (mean body mass = ~209 kg) and African-specific species in the dataset (mean body mass = ~166 kg). The 0.95 quantile displacement distances averaged roughly 13 km for all mammals in the full model, 11 km for the larger bodied mammals, 2 km for the smaller bodied mammals and 18 km for African mammals.

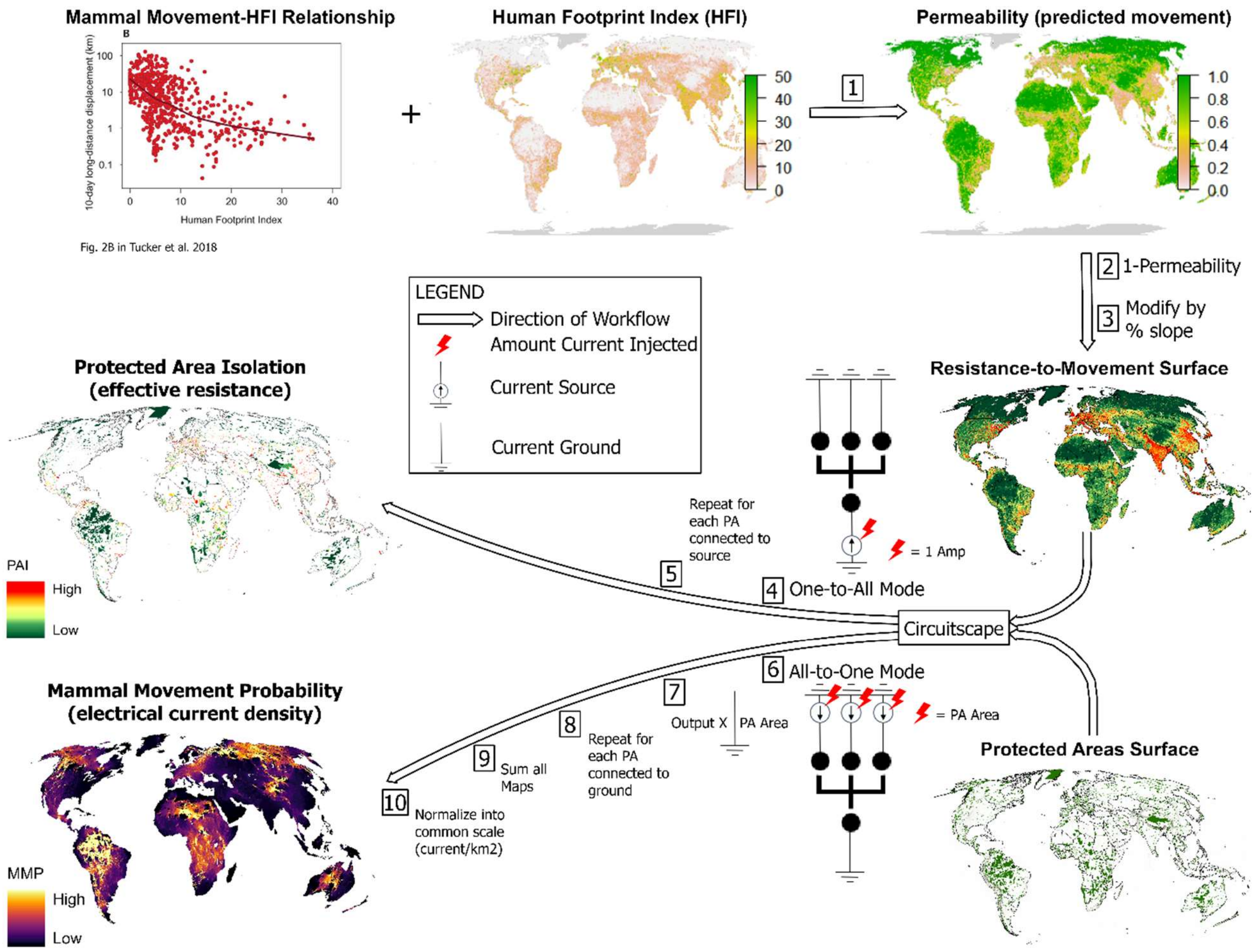


Fig. S10. Workflow diagram. Starting from upper left corner, moving to upper right corner following arrows to the two results: protected area isolation (PAI) and the global map of predicted mammal movement probability (MMP).

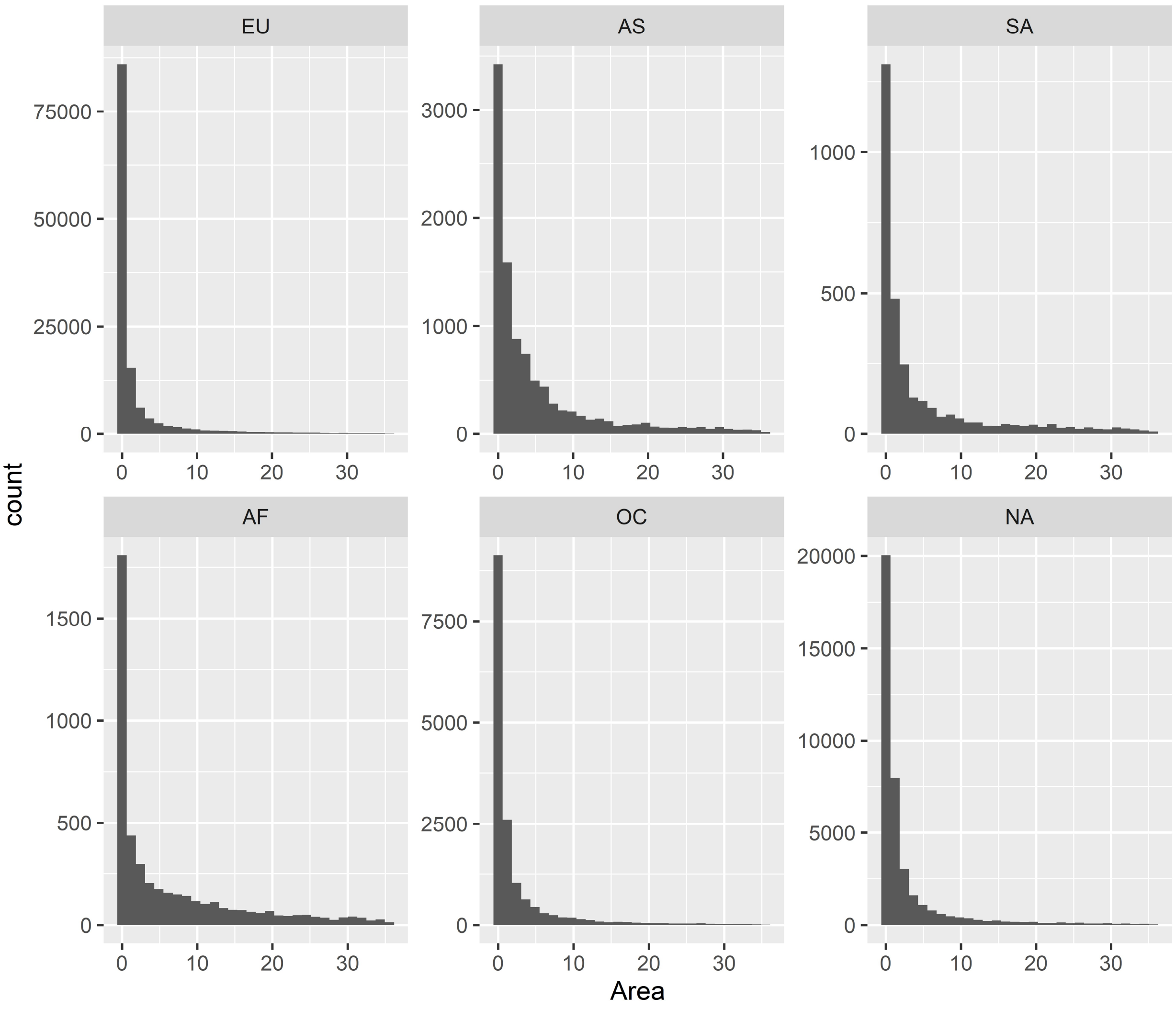


Fig. S11. Distribution of protected area (PA) sizes ($km^2$) that were excluded as sources and destinations of mammal movement. The size of excluded PAs ranged from < 0.1 to 35 $km^2$, with a mean of 2.4 $km^2$. Distributions are organized by continent: EU = Europe; AS = Asia; SA = South America; AF = Africa; OC = Oceania and Australia; NA = North America.

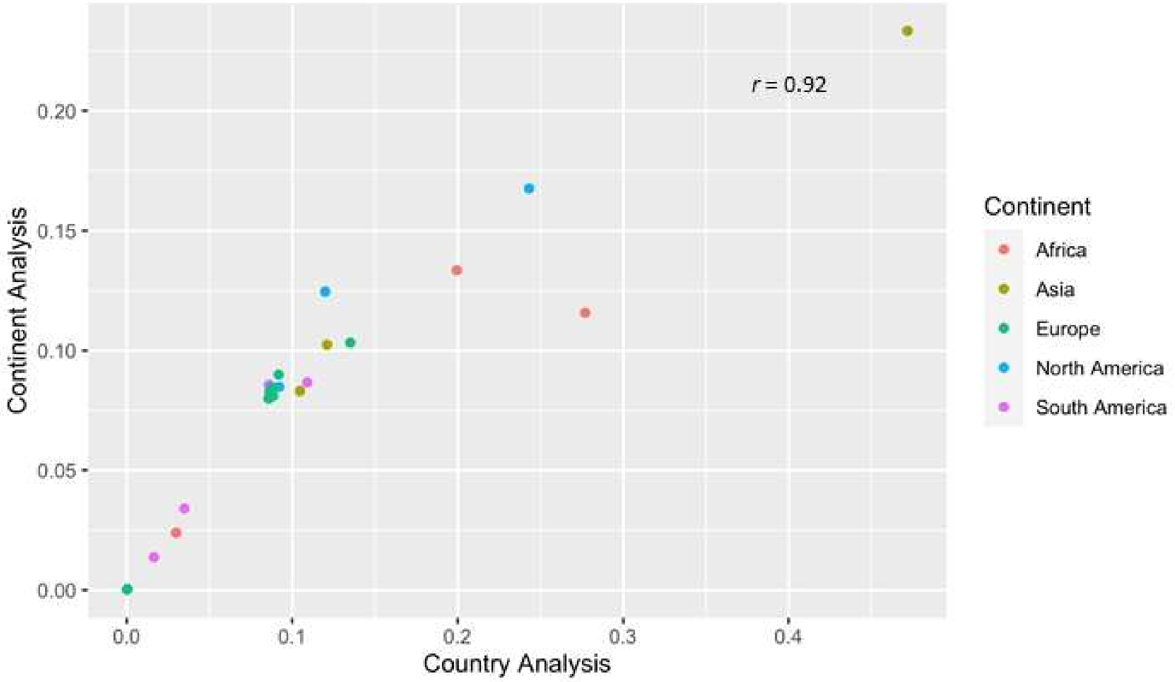


Fig. S12. Sensitivity of national protected area isolation (PAI) to international protected areas (PAs). Comparison of national PAI for 25 randomly selected countries estimated from the global analysis, conducted at the continent scale (y axis), and the country-scale analysis (x axis). The two measures of national PAI are highly correlated (Pearson's $r = 0.92$), indicating that measures taken from the global analysis are suitable proxies of national PAI.

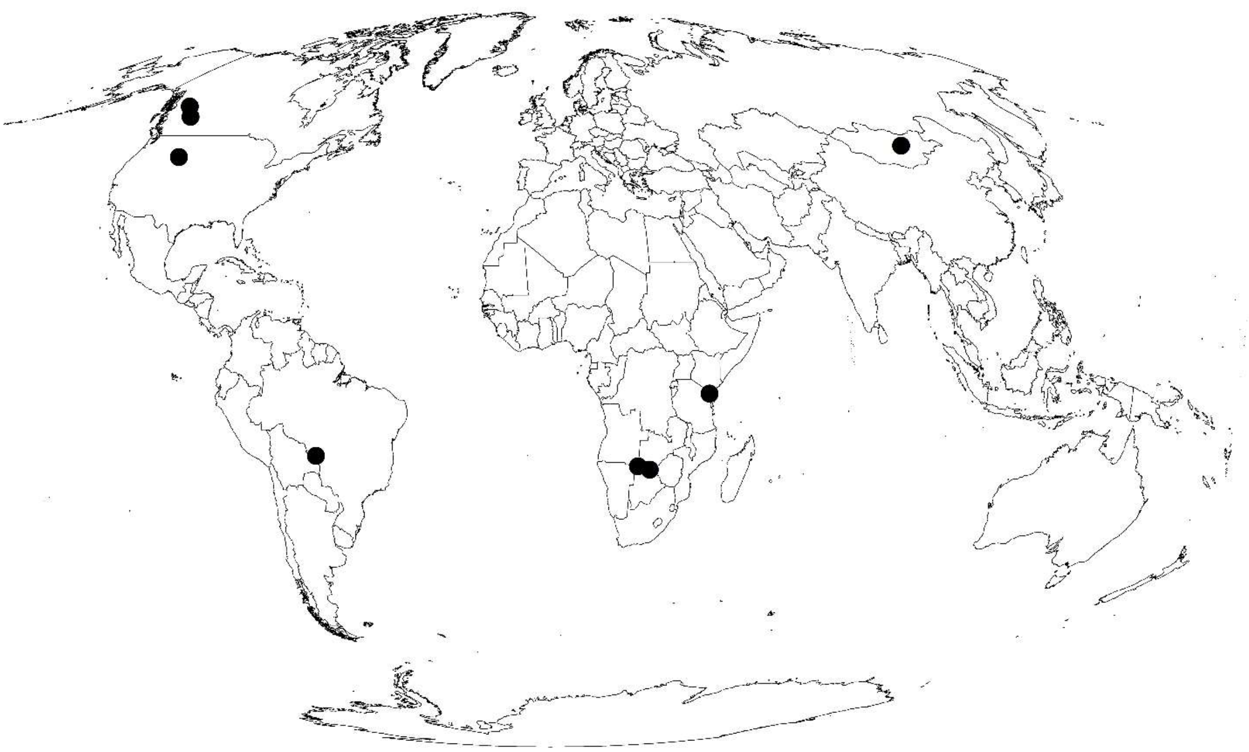

Fig. S13. Validation study locations. General locations of GPS datasets used to validate the global connectivity model. Datasets include more than 1.7 million GPS locations for 454 collared individuals from 11 species of mammals (see Table S1 for details and citations).

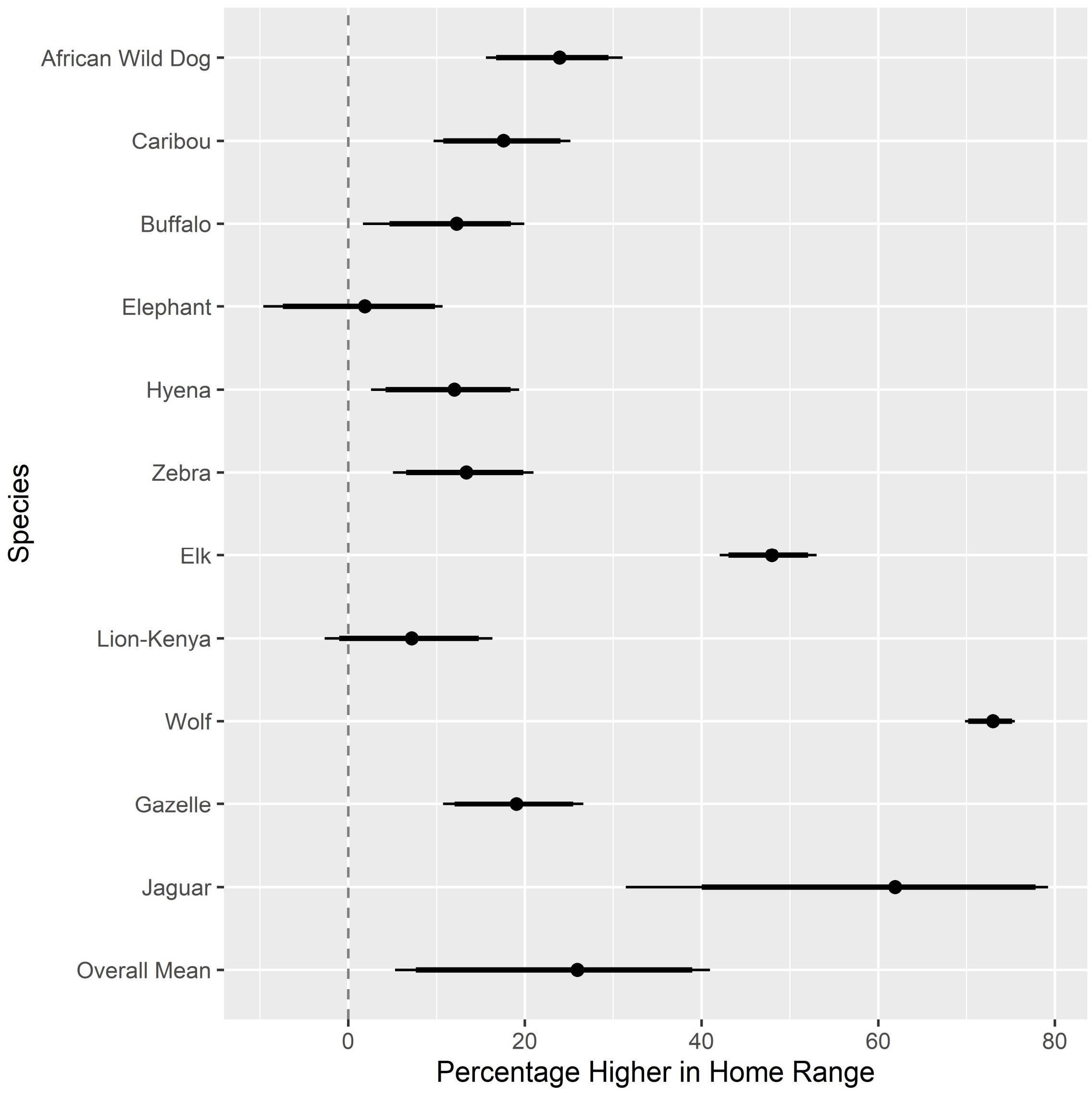


Fig. S14. Validation of global connectivity model. Estimated median difference in electrical current density (i.e., MMP) inside and outside estimated utilization distributions (UDs) estimated from each species GPS dataset. Dotted line = no difference inside or outside UDs; Thin black line = 95% CIs; Thick black line = 90% CIs.

Table S1. Top 5% connected countries for each connectivity indices and ranking for Canada and Russia.

| | PAI | Proportion of Protected Area* | ConnIntact* | PARC* |
|---|---|---|---|---|
| Top 5% Connected Countries | Greenland | New Caledonia | Greenland | Greenland |
| | Canada | Venezuela | Guyana | French Guiana |
| | Guyana | Slovenia | Svalbard & Jan Mayen Islands | Namibia |
| | Svalbard & Jan Mayen Islands | Bhutan | French Guiana | Venezuela |
| | French Guiana | Brunei Darussalam | Brunei Darussalam | Reunion |
| | Iceland | Greenland | Peru | Brazil |
| | Suriname | Luxembourg | Suriname | Botswana |
| | Finland | - | Brazil | Nicaragua |
| Rank for Canada** | 2 | 111 | 15 | 52 |
| Rank for Russsia** | 25 | 110 | 35 | 47 |

*Existing Global Indicators
**Two countries with highest proportion of intact wilderness.

Bold countries are those that overlap with our protected area isolation (PAI) metric.

Table S2. Summary of small PAs excluded from analysis.

| | Number of Used PAs | Number All PAs | Proportion of PAs Excluded | Area of Used PAs ($km^2$) | Total Area of All PAs ($km^2$) | Proportion of PA Area Excluded |
|---|---|---|---|---|---|---|
| Oceania* | 1343 | 17142 | 0.922 | 1638961 | 1677406 | 0.023 |
| South America | 2007 | 5080 | 0.605 | 5029290 | 5044512 | 0.003 |
| North America | 3871 | 42757 | 0.909 | 3885675 | 3990714 | 0.026 |
| Africa | 2627 | 7205 | 0.635 | 4788191 | 4811446 | 0.005 |
| Eurasia | 11243 | 146416 | 0.923 | 6022272 | 6299831 | 0.044 |

*Includes Oceania and Australia

Table S3. Summary of GPS movement data used to validate global connectivity model.

| Species | Location | Number Individuals | Number observations | Max Displacement (km) | Source* | Citations |
|---|---|---|---|---|---|---|
| African wild dog (*Lycaon pictus*) | Namibia | 14 | 19549 | 354 | 1 | Brennan et al.[5] |
| Caribou (*Rangifer tarandus*) | Canada | 186 | 245449 | 100 | 2 | Jones et al.[6]<br>Johnson et al.[7] |
| African buffalo (*Syncerous caffer*) | Namibia | 42 | 109941 | 128 | 1 | Brennan et al.[5] |
| African savanna elephant (*Loxodanta africana*) | Namibia | 73 | 1025411 | 273 | 1 | Brennan et al.[5] |
| Spotted Hyena (*Crocuta crocuta*) | Namibia | 20 | 32670 | 102 | 1,3 | Brennan et al.[5] |
| Plains zebra (*Equus quagga*) | Botswana | 14 | 26469 | 272 | 1,4 | Naidoo et al.[8] |
| Rocky Mountain elk (*Cervus canadensis*) | Wyoming | 17 | 104913 | 91 | 5,6 | Brennan et al.[9] |
| African lion** (*Panthera leo*) | Kenya | 3 | 2127 | 41 | 7 | NA |
| Grey wolf** (*Canis lupus*) | Canada | 68 | 174441 | 282 | 8 | Hebblewhite et al.[10]<br>Hebblewhite et al.[11] |
| Mongolian gazelle** (*Procapra gutturosa*) | Mongolia | 4 | 3308 | 257 | 9 | NA |
| Jaguar** (*Panthera onca*) | Brazil | 13 | 42909 | 34 | 10 | Morato et al.[12] |

*Sources
1. Provided by Namibian Ministry of Environment, Forestry and Tourism
2. Accessed from Movebank (movebank.org, study name "Mountain caribou in British Columbia", study ID 216040785; Movebank data repository DOI:10.544/001/1.p5bn656k)
3. Kwando Carnivore Project
4. WWF-US
5. Downloaded from Sciencebase (sciencebase.org)
6. Provided by Wyoming Department of Game and Fish, US Fish and Wildlife Service and US National Park Service
7. Accessed from Movebank (movebank.org, study name "Tsavo Lion Study", study ID 220229)
8. Accessed from Movebank (movebank.org, study name "Hebblewhite Alberta-BC Wolves", study ID 209824313)
9. Accessed from Movebank (movebank.org, study name "Mongolian Gazelle Mongolia WSCC", study ID 245174184); Provided by Wildlife Science and Conservation Center of Mongolia (WSCC)
10. Accessed from Movebank (movebank.org, study name "Movement ecology of the jaguar in the largest floodplain of the world, the Brazilian Pantanal", study ID 19411459)
** Not an independent dataset; i.e., used in Tucker et al.[4]

Table S4. Validation of critical connectivity area (CCA) use.

| Species | Percent that CCAs are used compared to the CCA available |
|---|---|
| African wild dog | *120.2* |
| Caribou | *113.0* |
| African buffalo | *142.8* |
| African savanna elephant | *147.6* |
| Spotted Hyena | *113.5* |
| Plains zebra | *10.0* |
| Rocky Mountain elk | *237.7* |
| African lion | *147.5* |
| Grey wolf | *257.5* |
| Mongolian gazelle | *0.0* |
| Jaguar | *148.6* |

Bold numbers indicate preference for CCA, i.e., percent used > 100
